# A role for ETV1 and endothelial cell-derived extracellular vesicle microRNAs in priming fibroblast response to vesicle-bound FGF2

**DOI:** 10.64898/2026.02.25.707568

**Authors:** Heidi Yuan, Chen Han, Lin Chen, Sriram Ravindran, Luisa A. DiPietro

**Affiliations:** Center for Wound Healing and Tissue Regeneration, University of Illinois Chicago, Chicago, Illinois 60612, USA; Department of Oral Biology, College of Dentistry, University of Illinois Chicago, Chicago, Illinois 60612, USA

**Keywords:** endothelial cells, wound healing, tissue regeneration, transcription factors, growth factors

## Abstract

Communication between various cell types following wounding is paramount for proper healing and regeneration of injured tissue. Endothelial cells and fibroblasts are critical cellular players involved in cutaneous wound repair, yet their communication mechanisms are not well understood. It has previously been shown that extracellular vesicles derived from endothelial cells (ECEVs) induce dermal fibroblasts to express a gene signature correlated with FGF2-mediated cancer associated fibroblast (CAF) activation, under the control of transcription factor ETV1. In this report, we utilize loss-of-function studies to define the mechanistic role of ETV1 in conferring this ECEV-induced transcriptomic shift and functional change in fibroblasts.

Additionally, we identify highly expressed ECEV microRNAs and examine their potential contribution to the ECEV mechanism through downstream gene modulation. In summary, we describe a plausible mechanism by which both ETV1 and top ECEV microRNAs promote a genotypic and phenotypic shift in dermal fibroblasts that have taken up ECEVs.

## Results & Discussion

Extracellular vesicles (EVs) are key mediators of paracrine signaling during wound repair (Narauskaitė et al. 2021). Endothelial cell-derived extracellular vesicles (ECEVs), released during angiogenesis (Todorova et al. 2017), likely coordinate communication between endothelial cells and neighboring fibroblasts during the proliferative phase of healing. We previously provided evidence to suggest that ECEVs regulate fibroblast proliferation and extracellular matrix (ECM) remodeling through vesicle-bound fibroblast growth factor 2 (FGF2) and the transcription factor ETV1 (Yuan et al. 2025), which regulates sensitivity to fibrotic signaling pathways among cancer-associated fibroblasts (Bordignon et al. 2019). Here, we define a mechanistic role for ETV1 in mediating ECEV-induced fibroblast responses and characterize ECEV microRNA (miRNA) cargo as a potential modifier of fibroblast signaling susceptibility.

We previously showed that ETV1 is upregulated in fibroblasts that have taken up ECEVs (Yuan et al. 2025). To define the contribution of ETV1 to ECEV mechanism, we assessed the impact of ETV1 loss-of-function on ECEV-induced fibroblast proliferation, one of the key phenotypes consistently enhanced in response to ECEV treatment. Transient transfection of ETV1 siRNA resulted in over 90% reduction of ETV1 mRNA within 48h (Figure 1a) and corresponding protein depletion by 72h (Figure 1b). Following transfection, we assessed fibroblast proliferation in response to ECEV treatment. Loss of ETV1 significantly attenuated ECEV-induced fibroblast proliferation, an effect not observed among control-transfected cells (Figure 1c), establishing ETV1 as an important mediator of fibroblast responsiveness to ECEVs and ECEV-bound FGF2.

**Figure 1.**
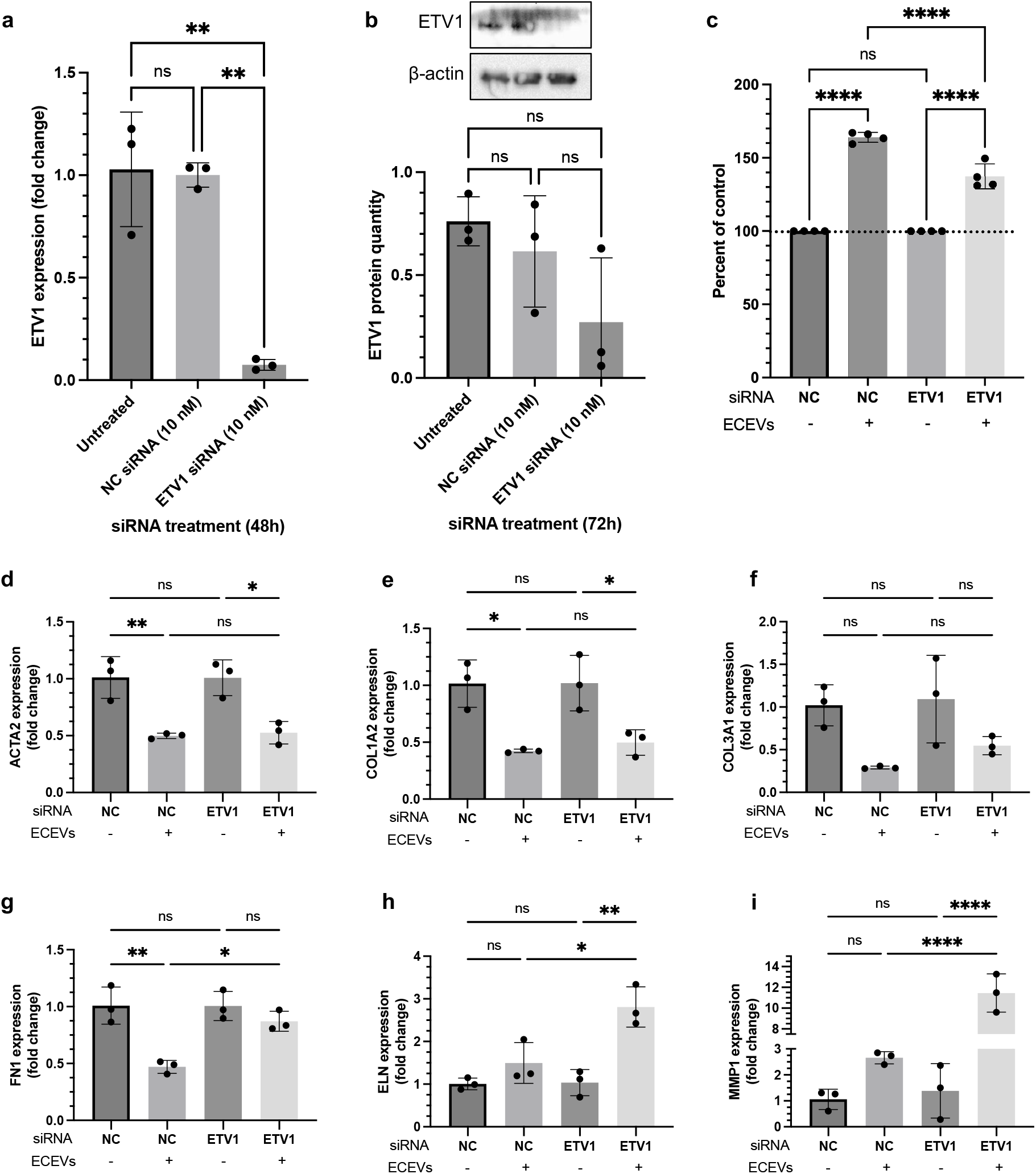
ETV1 knockdown attenuates ECEV induction of fibroblast proliferation and ECM gene dysregulation. (a) ETV1 gene expression levels in fibroblasts transfected with 10 nM of NC or ETV1 siRNA for 48h as determined by RT-PCR. (b) ETV1 protein levels were assessed by Western blot following 72h of transfection and normalized to β-actin. (c) CCK-8 assay was used to assess proliferation of NC or ETV1 siRNA-transfected fibroblasts treated with ECEVs for 48h (2 ECEV treatments, 24h apart). OD values are reported as a percentage of the untreated (no ECEV) fibroblasts within each siRNA group. (d) – (i) RT-PCR analysis of key ECM genes in NC or ETV1 siRNA-transfected fibroblasts subsequently treated with ECEVs for 24h. Data is presented as mean ± SD with data points indicating individual values from triplicate or quadruplicate cultures. Two independent experiments were performed and representative experiments are shown. One-way ANOVA with Bonferroni post-hoc test was used. ns = p > 0.05, * = p < 0.05, ** = p < 0.01, *** = p < 0.001, **** = p < 0.0001. ECEV, endothelial cell-derived extracellular vesicle; NC, negative control; OD, optical density.

ECEV-treated fibroblasts also exhibit a distinct ECM transcriptional profile characterized by downregulation of fibrotic and contractile genes (ACTA2, COL1A2, COL3A1, FN1, ELN) and upregulation of MMP1 (Yuan et al. 2025). These genes are regulated by ETV1 and associated with FGF2-mediated signaling (Bordignon et al. 2019). Following ETV1 knockdown, inhibition of ACTA2, COL1A3, COL3A1, and FN1 was partially reversed (Figure 1d-g), whereas changes in ELN and MMP1 expression were unaffected (Figure 1h-i). These findings indicate that ETV1 contributes selectively to ECM gene regulation downstream of ECEVs.

Beyond protein cargo, EVs transport miRNAs capable of modulating gene expression in recipient cells (Fernandes et al. 2022; Shirazi et al. 2021). Small RNA sequencing of ECEVs identified hsa-miR-126-3p, hsa-miR-151a-3p, hsa-miR-99a-5p, hsa-miR-21-5p, and hsa-miR-221-3p as the top 5 most abundant miRNAs (Figure 2a). To test whether miR-126-3p alone could recapitulate ECEV effects, fibroblasts were transfected with miR-126-3p mimics (Figure 2b). To our surprise, miR-126-3p overexpression led to a downregulation of proliferation and migration among fibroblasts (Figure 2c-d). Similar results were observed with mimics of the other top ECEV miRNAs (Figure S1), suggesting that individual miRNAs are insufficient to drive the proliferative effects of ECEVs.

**Figure 2.**
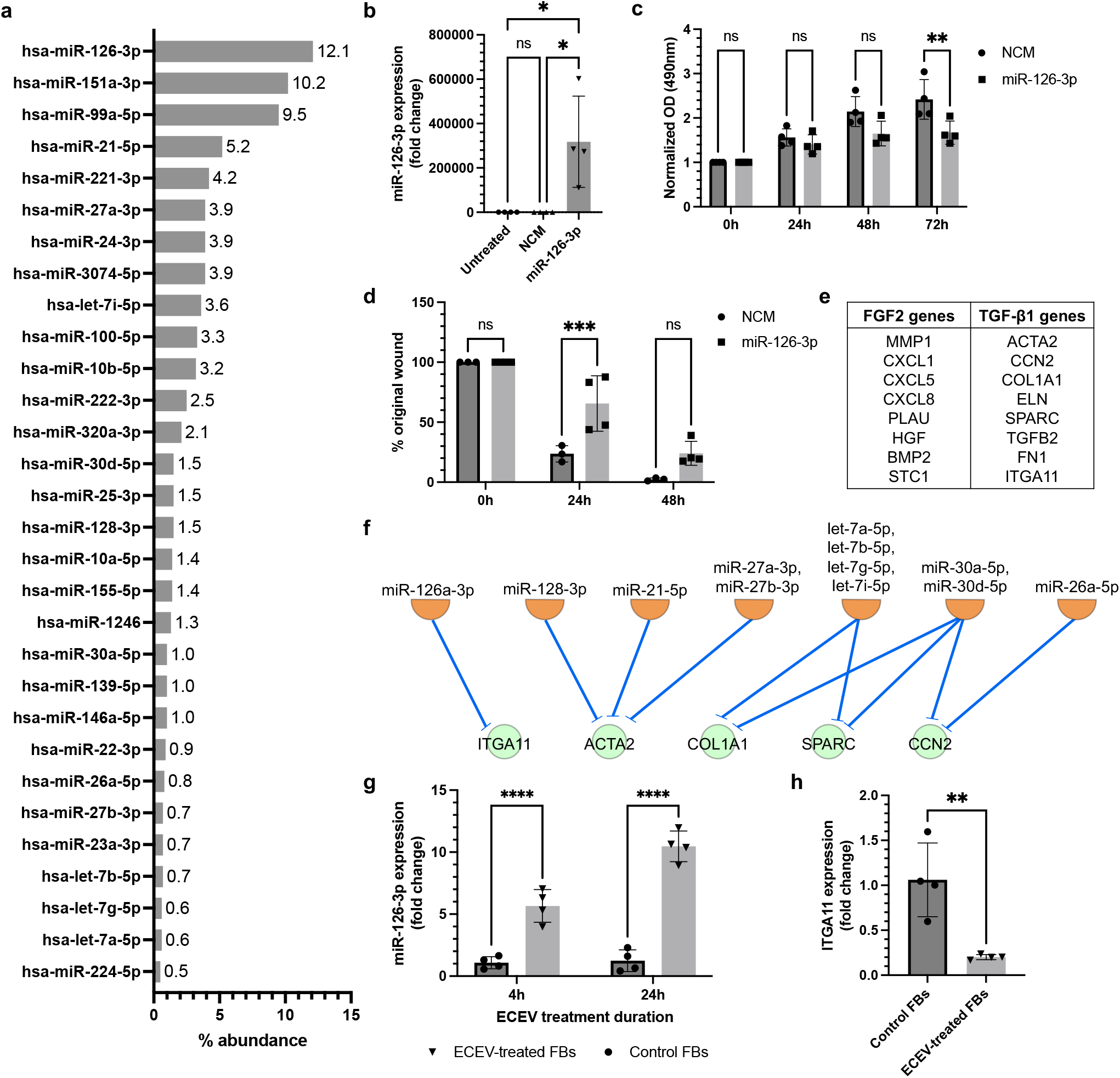
ECEV miRNAs repress TGF-β1-associated genes to enhance ETV1-mediated FGF2 signaling. (a) Top 30 highly expressed miRNAs found in ECEVs identified by small RNA sequencing. Data labels represent % abundance of the sequenced miRNA across total reads. (b) miR-126-3p expression was determined via RT-PCR following transfection of fibroblasts with mimics for 24h. (c) MTS assay was used to assess proliferation of miR-126-3p mimic-transfected fibroblasts. OD values were normalized to the initial cell density. (d) 3-well silicone inserts were used to assess migration of miR-126-3p mimic-transfected fibroblasts, with % original wound indicating percentage of the gap remaining at each time point. (e) FGF2-and TGF-β1-associated gene signatures derived from Bordignon et al. (2019). (f) IPA network demonstrating predicted inhibition of TGF-β1-associated transcriptional targets (green circular nodes) by highly expressed ECEV miRNAs (orange semicircle nodes). miRNAs with shared seed sequences were grouped into a singular node. (g) miR-126-3p expression levels in fibroblasts treated with ECEVs for 4h and 24h as determined by RT-PCR. (h) Expression levels of TGF-β1-associated and miR-126-3p gene target ITGA11 as determined by RT-PCR of fibroblasts treated with ECEVs for 24h. Data is presented as mean ± SD with data points indicating individual values from triplicate or quadruplicate cultures. Two independent experiments were performed and representative experiments are shown. One-way ANOVA with Bonferroni post-hoc test was used for (b), two-way ANOVA with Bonferroni post-hoc test was used for (c), (d), and (g), and unpaired t-test was used for (h). ns = p > 0.05, * = p < 0.05, ** = p < 0.01, *** = p < 0.001, **** = p < 0.0001. ECEV, endothelial cell-derived extracellular vesicle; FB, fibroblast; NCM, negative control mimic; OD, optical density.

We therefore hypothesized that ECEV miRNAs act cooperatively with vesicle-bound growth factors to bias fibroblast signaling. To test this theory, we performed an *in silico* analysis to identify predicted miRNA-mRNA relationships between top ECEV miRNAs and downregulated fibrotic gene targets in ECEV-treated fibroblasts, which exhibit a pro-FGF2 and anti-TGF-β1 genotype (Yuan et al. 2025). Using the miRNA target filter in Ingenuity Pathway Analysis (IPA), we determined whether the top 30 ECEV miRNAs have known targets within the TGF-β1-related genes previously reported by Bordignon and colleagues (Figure 2e) (Bhattacharyya et al. 2016; Bordignon et al. 2019). IPA revealed that 7 of the top 30 ECEV miRNAs are predicted to directly repress 5 out of 8 TGF-β1-associated genes (Figure 2f) previously linked to fibroblast activation. This pattern suggests a collective miRNA-mediated repression of TGF-β1-responsive genes, favoring an FGF2-inducible transcriptome.

To provide evidence of functional miRNA transfer, we analyzed levels of miR-126-3p in fibroblasts following ECEV treatment. By 4h and 24h following ECEV treatment, levels of miR-126-3p were significantly higher in ECEV-treated fibroblasts compared to controls, indicating efficient vesicle-mediated delivery (Figure 2g). This transfer was not detected among other top ECEV miRNA candidates (Figure S2). As additional proof of concept, we showed that levels of the TGF-β1-associated gene ITGA11, which is also a transcriptional target of miR-126-3p, is effectively downregulated following 24h of ECEV treatment (Figure 2h). Together, these findings support a functional role for transferred miRNAs in targeted transcriptional modulation that predisposes fibroblasts to FGF2 signaling.

The findings from this study implicate ETV1 as an important mediator of ECEV effects on dermal fibroblast proliferation and ECM gene dysregulation. ETV1 knockdown not only attenuated fibroblast proliferation stimulated by ECEVs, but also partially attenuated gene expression of downregulated ECM genes associated with TGF-β1 activation of fibroblasts. Small RNA sequencing revealed that miR-126-3p is the most highly expressed miRNA within ECEVs, and this miRNA is highly expressed in fibroblasts following ECEV treatment. Although the transfer of this miRNA alone is unlikely to contribute to the observed functional changes of ECEVs on fibroblasts, it may cooperate with additional ECEV miRNAs to preferentially repress TGF-β1-associated genes in order to facilitate ETV1 upregulation and FGF2-related gene transcription.

Several limitations should be noted. A stable transfection approach for ETV1 knockdown would enable analysis of functions such as ECM deposition that require longer incubation times in cell culture. Additionally, IPA-based miRNA-mRNA predictions are inherently inferential and require validation using high-resolution approaches such as CLIP-seq (Darnell 2010; Helwak and Tollervey 2016). Lastly, detection of miRNA transfer was most evident for miR-126-3p due to its high vesicular abundance and low baseline expression in fibroblasts; more sensitive methods may reveal transfer of additional miRNAs.

In summary, our findings support a two-pronged mechanism by which ECEVs modulate fibroblast behavior: FGF2-driven ETV1 activation promoting proliferation and selective ECM gene regulation, alongside coordinated miRNA-mediated repression of TGF-β1-associated genes that primes fibroblasts for FGF2 responsiveness.

## Materials and methods

### 1. ECEV isolation and collection

Primary human dermal microvascular endothelial cells (Lifeline Cell Technology #FC-0042) were cultured and grown to confluence in vascular cell basal media (ATCC #PCS-100-030) supplemented with growth factors and 5% fetal bovine serum (FBS) (ATCC #PCS-110-041). To collect ECEVs, confluent cells were washed twice with PBS and culture media was replaced with exosome-free media, comprised of vascular cell basal media supplemented with growth factors and 5% exosome-depleted FBS (Gibco #A2720803). After 48 hours, the conditioned media was collected and centrifuged at 300 x g for 5 min to remove residual live cell debris. The supernatant was collected and centrifuged again at 3000 x g for 15 min to remove apoptotic cells and debris. A polyethylene glycol (PEG)-based isolation reagent (4x) was added to the resulting supernatant according to established protocols (C.-C. Huang et al. 2024; Umar et al. 2024). The supernatant was then left to incubate overnight with rotation at 4°C. Following this, the supernatant was centrifuged at 1500 x g for 30 min at 4°C to pellet the ECEVs. For all *in vitro* assays, ECEV pellets were re-suspended in a volume based on the quantity of cells from which they were derived. In general, 1 µL volume was used per 1 x 10^4^ endothelial cells.

### 2. Small RNA sequencing and analysis of ECEVs

Total RNA from ECEVs that were collected and pelleted as described previously was isolated and purified using Trizol reagent (Invitrogen, Waltham, MA) and the RNA Clean & Concentrator-5 kit (Zymo Research, Tustin, CA, USA). RNA samples were submitted to Novogene (Novogene Corporation Inc., Sacramento, CA, USA) for quality control (QC), library preparation, sequencing, and quantitation as per the in-house standard for gene expression analysis. Library preparation and sequencing were only carried forward if the sample passed the QC step. In brief, adaptors were ligated to the 3’ and 5’ ends of small RNA. First strand cDNA was synthesized following hybridization with a reverse transcription primer. The double-stranded cDNA library was then generated through PCR enrichment. After purification and size selection, libraries with insertions between 18 and 40 base pairs were ready for sequencing on Illumina sequencing with SE50. The library was checked with Qubit and real-time PCR for quantification, and the bioanalyzer was used for size distribution detection. Quantified libraries were pooled and sequenced on Illumina platforms, according to the effective library concentration and data amount.

For analysis of ECEV microRNA content, sequenced raw data (in FASTQ format) was first processed for quality control. In this step, clean reads were obtained by removing sequences containing poly-N; poly-A,T,C, or G; 5’ adaptor contaminants; missing 3’ adaptors or the insert tag; and any additional low quality reads from the raw data. Additionally, Q20, Q30, and GC-content of the raw data were calculated, and a length range from the clean reads was chosen from the clean reads to perform downstream analysis. The small RNA tags were mapped to reference sequences by Bowtie (Langmead et al. 2009), with up to 1 mismatch to analyze their expression and distribution. Mapped small RNA tags were used to identify known miRNA with miRBase22.0. Modified software mirdeep2 (Friedländer et al. 2012) and srna-tools-cli were used to obtain potential miRNA sequences and draw secondary structures. Custom scripts were used to obtain miRNA counts. Tags originating from protein-coding genes, repeat sequences, rRNA, tRNA, snRNA, and snoRNA were removed by mapping small RNA tags to RepeatMasker or Rfam database. To ensure each small RNA sequence mapped to only a single annotation, the following priority rule was employed: known miRNA > rRNA > tRNA > snRNA > snoRNA > repeat > gene > NAT-siRNA > gene > novel miRNA > ta-siRNA. Quantification of miRNA expression levels were estimated by TPM (transcript per million) based on the following formula: Normalized expression = mapped read counts / Total reads x 1,000,000. To obtain relative percentage reads for top identified ECEV miRNAs, read counts for each identified miRNA were summed for each ECEV sample (n = 3) and then averaged to obtain total miRNA reads. The average read count for each miRNA was then divided by the total miRNA reads and multiplied by 100 to obtain a percent read value.

### 3. In vitro cell culture and transfection of fibroblasts

For all cell culture experiments, primary human dermal fibroblasts (Fisher Scientific #C0045C) were cultured in DMEM (Mediatech, Inc.) with 10% FBS (Gemini). During transfection, no antibiotics were used in the media. After transfection, and to monitor functional responses, 100 U/mL penicillin-streptomycin (Life Technologies) was included.

Transfection of fibroblasts with ETV1 siRNA or miRNA mimics was performed at 80% cell density. Lipofectamine (Invitrogen) was used as the transfection reagent according to manufacturer instructions and Opti-MEM (Gibco) was used to prepare all mimic- or siRNA-lipid complexes. All siRNAs and mimics used, including controls and final concentrations, are listed in Table S1.

To assess ETV1 knockdown at the gene and protein level, primary human dermal neonatal fibroblasts were seeded at a density of 7 x 10^5^ cells per well in 12-well plates for RT-PCR analysis and 2 x 10^6^ cells per well in 6-well plates for Western blot analysis. Following initial transfection with siRNA, cells were harvested at 48h for gene expression analysis and 72h for protein (Western blot) analysis.

To assess proliferation following ETV1 knockdown, fibroblasts were seeded at a density of 5 x 10^3^ cells per well in a 96-well plate and left to adhere overnight. Transfection was performed as described above. 48h following initial transfection, ECEV or control treatment was administered. For ECEV treatment, ECEVs were collected according to the methods described above and resuspended at a ratio of 30% v/v in exosome-free media prior to addition to fibroblasts. This treatment concentration was determined as an optimal saturation point for fibroblast uptake of ECEVs according to previously published data (Yuan et al. 2025). Exosome-free media, used as a control treatment, comprised of DMEM (Mediatech, Inc.) supplemented with 10% exosome-depleted FBS (ThermoFisher Scientific, #A2720803) and 100 U/mL penicillin-streptomycin (Life Technologies). Cell proliferation was assessed at 24h and 48h following ECEV treatment using a CCK-8 assay (Abcam). A second ECEV treatment was applied following the 24h CCK-8 assessment. Optical density at 460 nm was assessed using a plate reader spectrophotometer (Molecular Devices).

To assess ECM gene dysregulation following ETV1 knockdown, fibroblasts were seeded at a density of 5 x 10^4^ cells per well in a 24-well plate and left to adhere overnight. Transfection was performed as described above. 72h following initial transfection, ECEV or control treatment was administered. Cells were collected 24h later for RT-PCR analysis for a panel of ECM genes.

To assess the effect of miRNAs on fibroblast function, fibroblasts were seeded at a density of 5 x 10^4^ cells per well in a 12-well plate and transfection of cells with miRNA mimics was performed as described above. 24h following transfection, fibroblasts were loaded at a density of 4 x 10^3^ cells per well into a 96-well plate for proliferation assays and 1.5 x 10^4^ cells per well into 3-well silicone culture-inserts (Ibidi) for migration assays. Cells were left to adhere overnight prior to initial assessments of function. For proliferation, MTS reagent (Promega) was added at 0, 24, 48, and 72h and absorbance (optical density) at 490 nm was measured using a plate reader spectrophotometer (Molecular Devices). For migration, silicone inserts were removed to create gaps within a confluent bed of cells. Migration was assessed at 0, 24, and 48h by determining the area of the gap using ImageJ analysis. Percent of the original gap (“wound”) remaining was determined at each time point. To confirm transfection efficiency, cells were harvested 48h following transfection from the original 12-well plates and miRNA expression was determined via RT-PCR.

To assess the transfer of miR-126-3p to fibroblasts through ECEVs, fibroblasts were seeded at a density of 3 x 10^5^ in 12-well plates and left to adhere overnight. ECEV or control treatment was administered as described above. 4h and 24h following treatment, fibroblasts were harvested for RT-PCR analysis to assess miRNA levels or target gene expression.

### 4. RT-PCR analysis

To assess levels of ETV1, miRNA target gene ITGA11, or ECM genes, total RNA from transfected and/or ECEV-treated fibroblasts was isolated using Trizol-chloroform extraction and purified with the RNA Clean & Concentrator-5 kit (Zymo Research, Tustin, CA, USA). RNA concentrations obtained from the extraction protocol was determined using the Nanodrop 2000 system (Thermo Scientific). 1 µg of total RNA per sample was subsequently treated with DNAse (ThermoFisher Scientific, Waltham, MA) and converted to cDNA using a High-Capacity cDNA Reverse Transcription Kit (Invitrogen). Relative expression of each gene was determined via quantitative RT-PCR on the StepOnePlus RealTime PCR System (Applied Biosystems, Waltham, MA) using Power SYBR Green PCR Master Mix (Roche, Basel, Switzerland). The 2^-ΔΔCt^ was used to calculate relative expression of each gene. GAPDH was used as a reference gene. All primers are listed in Table S2.

To assess relative miRNA expression of ECEV-treated fibroblasts, total RNA from transfected or treated fibroblasts was extracted as described above. MiRNA concentrations were determined using the QubitTM microRNA Assay Kit (Invitrogen) and the QubitTM Flex Fluorometer (Invitrogen). 120 ng of total miRNA was then polyadenylated and reverse-transcribed to cDNA using the miRNA 1st-Strand cDNA Synthesis Kit (Agilent #600036).

Relative expression of miR-126-3p was determined via quantitative RT-PCR on the StepOnePlus RealTime PCR System (Applied Biosystems, Waltham, MA) using the QuantiTect SYBR Green RT-PCR kit (Qiagen) and a Universal Reverse Primer (Agilent #600037) as the downstream primer for all miRNA targets. The 2^-ΔΔCt^ was used to calculate relative expression of miR-126-3p, using RNU6B as a reference gene. All forward/upstream primers for miRNA targets are listed in Table S2.

### 5. Western blot analysis

To quantify intracellular levels of ETV1, whole cell lysate from ETV1 siRNA-transfected fibroblasts was extracted using RIPA buffer (Sigma-Aldrich #R0278) containing 1% protease inhibitor cocktail (Sigma #P8340) followed by sonication and centrifugation at 16,000 x g for 15 min at 4°C to remove cell debris. Supernatant was collected and a BCA protein assay was performed (Pierce BCA #23225, Thermo Scientific) to determine total protein concentrations. 40 µg of total protein for each sample was then resuspended in Laemmli buffer (Bio-rad Laboratories) and β-mercaptoethanol as a reducing agent, then placed on a heat block at 95°C for 5 min. After cooling on ice, samples were loaded onto precast SDS-PAGE gels (Bio-Rad Laboratories), followed by transfer to PVDF membranes, then blocked with 5% non-fat milk in TBST buffer (Bio-Rad Laboratories) for 1 hour at room temperature. Following washing, the membranes were incubated with primary antibodies overnight at 4°C with rotation at the indicated dilutions: ETV1 (1:500, GeneTex, #GTX129202) and β-actin (1:1000, Cell Signaling, #5125). Membranes were washed and subsequently incubated in HRP-conjugated secondary antibody (1:2000, Bio-Rad Laboratories, #1706515) for 1 hour at room temperature. ECL substrate (Bio-Rad Laboratories) was added and chemiluminescence was detected using the ChemiDoc XRS Scanner (Bio-Rad Laboratories). Density analysis was performed using ImageJ’s Gel Analyzer function.

### 6. Ingenuity Pathway Analysis (IPA)

Ingenuity Pathway Analysis (IPA, Qiagen) was used to identify a subset of predicted fibroblast gene targets of the top ECEV miRNAs obtained from small RNA sequencing. Using the miRNA-mRNA target filter, a list of all predicted gene targets of the top 30 miRNAs (identified via TargetScan, miRecords, or Ingenuity Expert Findings), was filtered against 8 downregulated gene targets in ECEV-treated fibroblasts that overlap with TGF-β1-associated CAF effector genes (Figure 2e). Predicted inhibitory miRNA-mRNA relationships were displayed as a hierarchical network and reorganized based on most highly expressed to least highly expressed miRNA.

### 7. Statistical analysis

Data were expressed as mean ± standard deviation (SD) unless specified otherwise, and their normality was evaluated using the Kolmogorov-Smirnov or Shapiro-Wilk tests. Statistical comparisons were performed using a two-tailed unpaired t-test, a one-way ANOVA, or a two-way ANOVA followed by the Bonferroni post-hoc test using GraphPad Prism version 10.6.1 (GraphPad, San Diego, CA). P-values less than 0.05 were considered statistically significant.

## Ethics Statement

This study was performed in accordance with the Declaration of Helsinki. Human primary cell lines included in this study were approved as part of this study protocol. Ethical approval was not received for this human study because all primary cells were purchased from Lifeline Cell Technology or Fisher Scientific, which states that all donated tissues have been obtained under proper informed consent and adheres to the Declaration of Helsinki, The Human Tissue Act (UK), CFR Title 21, and HIPAA Regulations relative to obtaining and handling human tissue for Research Use.

## Data Availability Statement

Datasets related to this article can be found in NCBI GEO with accession number GSE316721. Primary data related to this publication can be accessed via INDIGO with the DOI: https://doi.org/10.25417/uic.31151866.

## Acknowledgments

This research was funded by NIH R01-GM50875 & R35-GM139603 (LAD), F31-AR083830 (HY), F31-AR082287 (CH), R01-DE027404 & DE30495 (SR).

## Author Contributions

Conceptualization: HY, SR, LAD; Data curation: HY, CH, LC; Formal analysis: HY, CH, LC, SR, LAD; Funding acquisition: HY, LAD, CH, SR; Investigation: HY, CH, LC; Methodology: HY, CH, LC, SR, LAD; Visualization: HY; Writing – original draft: HY, LAD; Writing – review & editing: HY, CH, LAD, SR, LC

---

**Figure S1.**
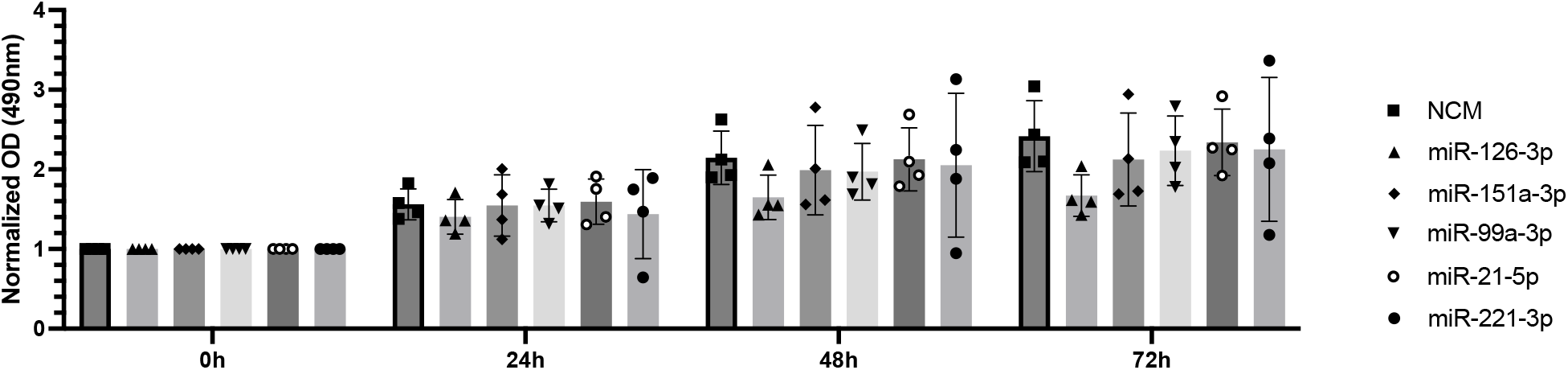
Transfection of dermal fibroblasts with mimics of top 5 ECEV miRNAs fails to re-capitulate proliferative fibroblast phenotype induced by ECEVs. MTS assay was performed on dermal fibroblasts transfected with mimics of the top 5 indicated ECEV miRNAs to assess proliferation. OD values were normalized to the initial cell density. Data is presented as mean ± SD with data points indicating individual values from quadruplicate cultures. Two-way ANOVA with Bonferroni post-hoc test was used and no significant differences between groups were found. NCM, negative control mimic; OD, optical density.

**Figure S2.**
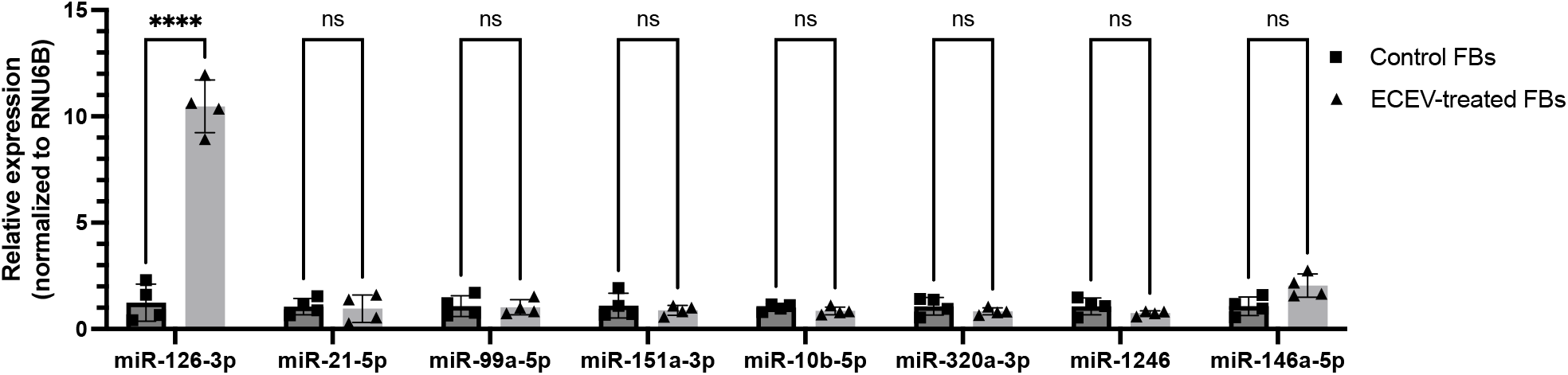
ECEV treatment of fibroblasts leads to detectable transfer of miR-126-3p but not other top ECEV miRNAs. Fibroblasts treated with ECEVs for 24h were analyzed for miRNA expression analysis via RT-PCR. Relative expression is normalized to RNU6B. Data is presented as mean ± SD with data points indicating individual values from quadruplicate cultures. Two-way ANOVA with Bonferroni post-hoc test was used. ns = p > 0.05, * = p < 0.05, ** = p < 0.01, *** = p < 0.001, **** = p < 0.0001. ECEV, endothelial cell-derived extracellular vesicle; FB, fibroblast.

**Table S1.** siRNA and miRNA mimics used for transfection.

| siRNA or mimic | Assay ID | Catalog # | Source | Final concentration |
| --- | --- | --- | --- | --- |
| Silencer™ Cy™3 Labeled Negative Control #1 siRNA |  | AM4621 | Thermo Fisher Scientific | 10 nM |
| ETV1 Silencer® siRNA | 115583 | AM16708 | Thermo Fisher Scientific | 10 nM |
| Negative Control #1 miRVana™ miRNA mimic |  | 4464058 | Thermo Fisher Scientific | 20 nM |
| hsa-miR-126-3p mirVana® miRNA mimic | MC12841 | 4464066 | Thermo Fisher Scientific | 20 nM |
| hsa-miR-151a-3p mirVana® miRNA mimic | MC11405 | 4464066 | Thermo Fisher Scientific | 20 nM |
| hsa-miR-99a-3p mirVana® miRNA mimic | MC10719 | 4464066 | Thermo Fisher Scientific | 20 nM |
| hsa-miR-21-5p mirVana® miRNA mimic | MC10206 | 4464066 | Thermo Fisher Scientific | 20 nM |
| hsa-miR-221-3p mirVana® miRNA mimic | MC10337 | 4464066 | Thermo Fisher Scientific | 20 nM |

**Table S2.**
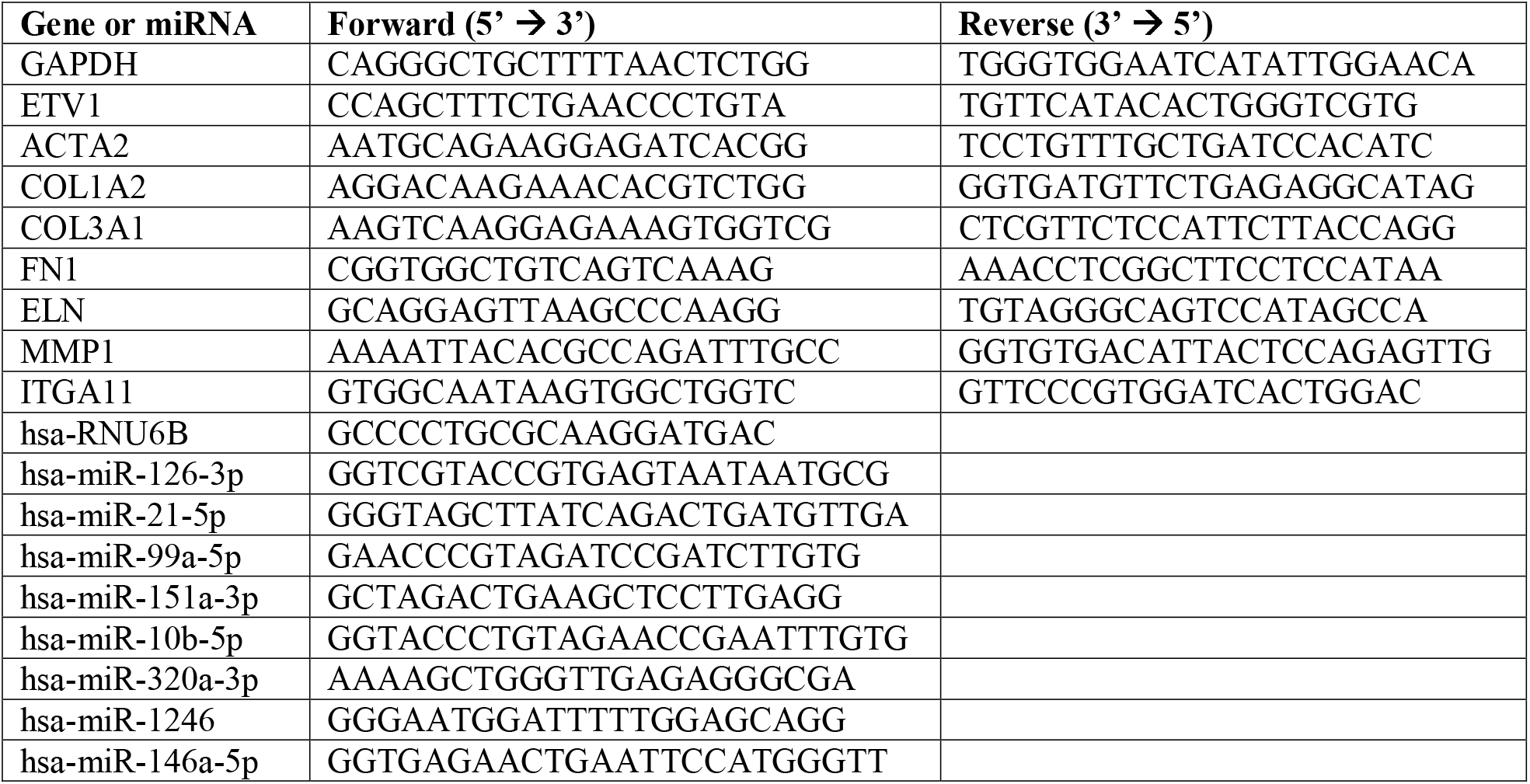
Primer sequences for fibroblast gene and miRNA expression analysis.

